# An ELISA for discovering protein-protein interaction inhibitors: blocking lysinoalanine crosslinking between subunits of the spirochete flagellar hook as a test case

**DOI:** 10.1101/2025.10.31.685777

**Authors:** Maithili Deshpande, Michael J. Lynch, Kurni Kurniyati, Chunhao Li, Brian R. Crane

## Abstract

The inhibition of a specific protein-protein interaction is often difficult to achieve in targeted drug design. We report the development and optimization of a general-purpose, readily implemented enzyme-linked immunosorbent assay (ELISA) for high-throughput screening to identify small-molecule inhibitors of protein interactions. This ELISA does not involve the use of any capture antibodies, probes, or compounds coated on the plate and represents a general strategy to identify inhibitors of a given protein-protein interaction. We demonstrate its utility in blocking lysinoalanine crosslinking between subunits of the spirochete flagellar hook by targeting the native form of the FlgE protein, which differs from the strategies used in previous assays. The flagellar hook protein FlgE self-catalyzes the formation of a lysinoalanine (Lal) inter-subunit crosslink that is essential for the motility, and thus, infectivity of spirochetes. Prevention of Lal crosslinking through inhibition with small molecules thus represents an avenue for therapeutic development against spirochete-related diseases, such as Lyme and syphilis. Screening a library of ∼700 compounds with the ELISA confirmed that hexachlorophene, currently the only known inhibitor of Lal crosslinking in FlgE, effectively inhibits the crosslinking reaction. In addition, the assay identified two new potential inhibitors, honokiol and zafirlukast, and several activators which belong to well-known classes of antibiotics.

## Introduction

Protein-protein interactions (PPIs) underlie much of metabolism. Their functions include: enzyme regulation, modulating protein activity, transport and trafficking of molecules, host-pathogen interactions, signal transduction between cells, cell-to-cell interactions, and metabolic and developmental control^1–3^. Rapid, inexpensive assays that could accelerate inhibitor discovery to therapeutically target PPIs are highly desired in the drug discovery field. This study focuses on the development and optimization of a heterogeneous enzyme-linked immunosorbent assay (ELISA), which does not involve the use of capture antibodies or probes coated on the plate, as a general strategy to identify inhibitors of a PPI. It was developed alongside homogeneous or separation-free immunoassays in which measurements are performed in solution without prior separation of free and bound components^4,5^ in efforts to identify antimicrobial compounds against spirochete bacteria.

Spirochetes cause several chronic and stage-related diseases including syphilis (*Treponema pallidum* ssp *pallidum* [Tps]), Lyme disease (*Borrelia burgdorferi* [Bb]), and periodontal disease (*Treponema denticola* [Td])^6–13^. In recent years, the emergence of multi drug-resistant strains to commonly used antibiotics (e.g., β-lactams, erythromycin, tetracyclines, azithromycin, rifampin, etc.)^14,15^ and the remarkable structural and physiological variability of spirochetes compared to other bacterial pathogens^16,17^ have refocused efforts to develop novel antimicrobials and therapeutic strategies^5,18^. We have focused on targeting a covalent post-translational modification, the lysinoalanine (Lal) crosslink, in the spirochete bacterial flagellar hook protein FlgE^19,20^. FlgE self-catalyzes the formation of the Lal covalent linkage between conserved lysine and cysteine residues of two adjacent FlgE subunits^19,20^, which polymerizes the hook into a stable high-molecular-weight complex (HMWC)^16,21,22^. Lal formation involves conversion of a Cys178 residue to a dehydroalanine (DHA) intermediate, followed by the reaction of DHA with a specific Lys165 of another FlgE molecule to produce Lal (LA165:178 in *Td* FlgE). The prevention of Lal crosslinking through mutation and its inhibition with small molecules, impairs motility in spirochetes^19^. As motility is essential for the infectivity and invasiveness of spirochetes^16,17,23–27^, inhibiting the formation of the Lal crosslink with small molecules provides a viable method to treat spirochete infections.

Our initial efforts at targeting the Lal crosslink were focused on a homogeneous luminescence-based assay^5^. This assay utilizes the complementation of a split-luciferase enzyme (NanoLuc) by the covalent association of two engineered and chemically primed FlgE domains. Although this assay has good signal-to-noise ratio and dynamic range, inhibition manifests whether a compound prevents Lal formation or assembly of the split luciferase units. Moreover, the assay uses FlgE in which Cys is pre-converted to the intermediate DHA, and thus biases for inhibitors that recognize the DHA-harboring conformation, which differs considerably from the native Cys-containing protein^20^. DHA conversion not only eliminates sulfide, but also rearranges the surrounding secondary structure.

As homogeneous assay technology does not involve separation steps, it is commonly used in high throughput screening (HTS) of compound libraries. However, heterogeneous assays such as the conventional enzyme-linked immunosorbent assay (ELISA) have their own advantages. Automated ELISA workflow stations use automated liquid handlers for sample preparation, reagent addition and washing steps, and robotic arms that can handle plate transfers^28^. Heterogeneous assays are often more precise and subject to less interference than homogeneous assays, thus benefiting the detection of more complex analytes^29–31^. Antibody-based ELISA systems have been routinely used to detect large protein molecules in cell lysates and in clinical diagnostics for marker proteins^32,33^. However, to our knowledge, ELISAs for the direct detection of PPIs between homomeric or oligomeric proteins that use the passive adsorption of one of the protein interacting partners on the plate without the involvement of primary capture antibodies, probes, or compounds, are rare.

We herein report the development and optimization of an ELISA assay for inhibitors of PPIs. This modified sandwich assay, in which the FlgE protein is conjugated with a fluorescein isocyanate (FITC) ethylenediamine derivative, and PPIs are detected with an anti-FITC:horseradish peroxidase (HRP) conjugate, has high sensitivity and relatively low detection times. Unlike the split-luciferase NanoLuc complementation assay^5^, the ELISA assay is based on native FlgE protein and hence can in principle target both the pre-reactive Cys state as well as the DHA intermediate state. The method has rapid read-out, even though the chemistry of Lal crosslinking is slow. To test the applicability of this assay, we validated it with a known Lal inhibitor, and in a small library screen, discovered several new compounds, one of which has activity in *T. denticola* cell culture.

## Materials and Methods

### Protein expression and purification

*Treponema denticola* (Td) FlgE G11-M454 and its corresponding C178A variant in plasmid pET28a containing a 6x His tag was recombinantly expressed in *E. coli* BL21 (DE3) cells^4^. Cells were grown in Luria-Bertani (LB) broth containing 50 µg/mL kanamycin at 37°C to an OD600 of ∼0.8 and induced with 0.5 mM IPTG at 18°C overnight. Cells were harvested after ∼16 h by centrifugation at 4000 x g for 15 min at 4°C. Cells were resuspended in lysis buffer (25 mM Tris pH 7.5, 500 mM NaCl, 30 mM imidazole, 10% glycerol) and lysed using an Avanti Emulsiflex C3 high pressure homogenizer. The lysate was centrifuged at 75000 x g for 1 h at 4°C and the supernatant was incubated with Ni-NTA resin for 1 h. The resin was washed with 10 column volumes (CV) of lysis buffer followed by 5 CV of wash buffer (25 mM HEPES pH 7.5, 500 mM NaCl, 30 mM imidazole, 10% glycerol). Protein was eluted in elution buffer (25 mM HEPES pH 7.5, 500 mM NaCl, 300 mM imidazole, 5% glycerol), concentrated in a 10 kDa MWCO spin concentrator (Amicon) and injected onto an S200 26/60 gel filtration (GF) column pre-equilibrated with GF buffer (25 mM HEPES pH 7.5, 150 mM NaCl). Fractions containing the protein were pooled after confirmation of protein purity by SDS-PAGE. The protein was either used to prepare the FlgE:5-FITC ethylenediamine conjugate or flash-frozen in liquid nitrogen and stored at -80°C to use as coat protein in the modified ELISA screening assay.

### Preparation of the FlgE:5-FITC ethylenediamine conjugate

The FlgE:5-FITC ethylene diamine conjugate was prepared by reacting WT *Td* FlgE G11-M454 or the C178A variant at 2 mg/ml in GF buffer with 10x molar excess of 5-FITC ethylenediamine dissolved in DMSO in the presence of 25x molar excess of N-hydroxysuccinimide (NHS) and 1- Ethyl-3-(3-dimethylaminopropyl)carbodiimide (EDC) in the dark for 2 h at room temperature. The conjugated protein was then desalted using BioRad Econo-Pac 10DG desalt columns equilibrated in GF buffer, aliquoted in small volumes, flash frozen in liquid nitrogen and stored at -80°C.

### Fluorescence measurement of the FlgE:5-FITC ethylenediamine conjugate

To verify conjugation of 5-FITC ethylenediamine to FlgE, 30 μL of 20 μM of FlgE:fluorophore conjugate was added to wells in a Corning black polystyrene, 96-well half-area plate. The sample was excited at 470 nm to prevent overlap between excitation and emission spectra, with cutoff at 495 nm, on a SpectraMax M5 microplate reader. Emission spectra were collected from 410-700 nm. GF buffer and non-conjugated *Td* FlgE were used as controls.

### Development of the ELISA

The concentrations of reagents, type of blocking solution used, temperature at which the assay was performed, antibody dilution used, and time of color development were optimized to generate the assay protocol as follows: either Nunc MaxiSorp modular strips or 96-well plates with flat-bottom wells (ThermoFisher) were used to perform the assays. The strips were coated with FlgE (100 μg/ml in PBS, pH 7.2; 100 μL/well) for 18-20 h at 4°C. The first well in each strip was uncoated for use as an assay blank. After coating, the strips were washed 6x (200 μL/well) with PBS containing 0.05% Tween-20 (PBST) and then blocked with 2% non-fat dry milk (NFDM) (200 μL/well for 4 h at room temperature). After 5x washes, the FlgE:5-FITC ethylenediamine conjugate (10 μg/mL, 100 μL/well) in crosslinking buffer (40 mM Tris pH 8.5, 160 mM NaCl, 1 M ammonium sulfate) was added to the wells. N-ethyl maleimide (NEM) at a final concentration of 1 mM was added to the wells and the Lal crosslinking reaction was allowed to proceed for 18- 24 h at room temperature. The wells were then washed 6x with PBST, strips were incubated with anti FITC:HRP conjugate (Invitrogen, catalog number PA1-86159, RRID identifier AB_930865) (1:15000 dilution in PBS, 100 μL/well) and shaken at 320 rpm for 1 h at room temperature. ABTS [2,2’-azino-bis(3-ethylbenzothiazoline-6-sulfonic acid] reagent was freshly prepared by dissolving ABTS to a final concentration of 1 mM in 70 mM citrate-phosphate buffer, pH 4.2 (147 mL of 0.1 M anhydrous citric acid + 103 mL of 0.2 M dibasic sodium phosphate) and 1 μL of 30% H2O2 was added per ml of ABTS solution immediately prior to use. After 6x washes, ABTS substrate solution was added, and the enzymatic reaction was stopped after 15 min by adding 1% SDS. Absorbance was measured at 405 nm.

For testing the inhibitors, the following modification was made. A 50 μL/well aliquot of an appropriate concentration of the test compound was added to each well followed by the addition of 50 μL/well of the FlgE:5-FITC ethylenediamine conjugate (20 μg/mL). The remaining procedure was the same.

The Z’ values for the assay were calculated using the formula Z’ = 1 – 3(𝜎_!_ + 𝜎_"_)/|𝜇_!_ −𝜇_"_| where (𝜎_!_ + 𝜎_"_) is the sum of the standard deviations of the positive and negative controls and |𝜇_!_ − 𝜇_"_| is the absolute value of the difference in means of the positive and negative controls.

### HTS of NIH clinical compound library

A library of 727 clinically-approved compounds was obtained from the National Institutes of Health (NIH), Bethesda, MD. ELISAs were performed as described above for testing the inhibitors. A cut-off value of 45% inhibition of the Lal crosslinking, as compared to the control, was used to identify potential lead compounds in the ELISA assay and further tested in the electrophoresis assay. The Z’ values for the assays were calculated as described above on 5 replicates each of the positive and negative control. Wherever possible, the data were also compared to that reported earlier using the NanoLuc assay^5^.

### SDS-PAGE Lal crosslinking assay

Compounds that gave ≥45% inhibition in the ELISA assay were further tested using SDS-PAGE as a confirmatory assay. Samples were prepared using 20 μM of *Td* FlgE G11-M454 in crosslinking buffer. Test compound and then NEM were added to a final concentration of 500 μM and 1 mM respectively. The crosslinking reaction was allowed to proceed for 24 h at 4°C. The reaction was quenched with 4x Laemmli reducing sample buffer and the samples were loaded on a 4-20% Tris-Glycine gel. Gels were stained with Coomassie blue for 1 h and destained to image the protein bands. Densities of the higher molecular weight complexes formed on Lal crosslinking were analyzed using ImageJ software.

### IC50 determination of inhibitor compounds

The ELISA assay was performed on a dilution series of the top compound hits obtained after the ELISA HTS and the secondary SDS-PAGE confirmatory assay. The IC50 values were obtained by fitting the results to the four-parametric log(inhibitor concentration) vs response equation curve Y = Bottom + (Top - Bottom)/(1 + 10^((LogIC50 - X)*HillSlope)) in GraphPad Prism, where X is log(inhibitor concentration) and Y is the percentage of inhibition compared to no-compound control.

### *T. denticola* cell culture and FlgE western blot analysis

*Treponema denticola* ATCC 35405 (wild-type) was used in this study. Cells were grown in tryptone-yeast extract-gelatin-volatile fatty acids-serum (TYGVS) medium at 37°C in an anaerobic chamber in the presence of 85% nitrogen, 5% carbon dioxide, and 5% hydrogen. For the zafirlukast-treated cells, a total of 10⁵ cells/mL of late-log phase *T. denticola* cultures was inoculated into 20 mL of TYGVS medium. Zafirlukast (100 µL of a 10 mM solution in DMSO, yielding a final concentration of 50 µM) was added to the culture, while 100 µL of DMSO was added as a control. The cultures were grown to late-log phase and subsequently used for further experiments. For the hexachlorophene-treated cells, a total of 10⁵ cells/mL of late-log phase *T. denticola* cultures was inoculated into 3 mL of TYGVS medium. Hexachlorophene (5 µL of a 50 mM solution in DMSO, yielding a final concentration of 83.3 µM) was added to the culture, while 5 µL of DMSO was added as a control. Cells were grown to the stationary phase and subsequently passaged for three additional generations in the presence of either hexachlorophene or DMSO.

Cultures were then plated on TYGVS plates containing 0.75% SeaPlaque agarose (Lonza, Durham, North Carolina) with either hexachlorophene or DMSO. For western blots, *T. denticola* cells were harvested at the late-log phase (∼10^9^ cells/ml). Cells were lysed via sonication and protein concentration determined via BCA assay. Samples were normalized to total protein concentration, mixed 1:1 with 2x Laemmli buffer supplemented with excess BME and heated at 95 °C. Samples were then centrifuged and loaded on an SDS-PAGE gel and electrophoresed at 200V in Tris-glycine SDS running buffer. The gels were rinsed with transfer buffer (25 mM Tris, 192 mM glycine, pH 8.3) and transferred to a 0.2 um PVDF membrane. Membranes were then blocked in 5% (w/v) skim milk in tris-buffered saline supplemented with 0.2% (v/v) Tween20 (TBST) for one hour at room temperature and incubated with a 1:1000 (v/v) dilution of FlgE antibody (αFlgE) in 5% (w/v) skim milk/TBST for one hour at room temperature. Membranes were then washed three times with TBST, incubated with 1:5000 (v/v) dilution of αRat HRP-IgG in 5% (w/v) skim milk/TBST for one hour and washed three times with TBST. Chemiluminescent bands were visualized via the addition of HRP chemiluminescent substrate and imaged within 30 seconds.

### *T. denticola* swimming plate assays

Swimming plate assays were performed with plates containing 50% (v/v) TYGVS:50% (v/v) PBS with 0.35% (w/v) SeaPlaque agarose. For this assay, a total of 10⁵ cells/mL of late-log phase *T. denticola* cultures was inoculated into 5 mL of TYGVS medium containing zafirlukast (50 µM) or hexachlorophene (83.3 µM) or DMSO alone (control) and grown anaerobically at 37°C until the cell density reached approximately 5x10^8^ – 10^9^ cells/mL. The resulting cultures were centrifuged and then resuspended in ∼25 μL of spent culture supernatant. Each plate was inoculated with 4 μL of resuspended cell cultures and incubated anaerobically at 37°C for 3 days to allow the cells to swim out. Two non-motile mutants, *Δtap1* (a*fliK* deletion mutant) or *ΔflgE* (a *flgE* deletion mutant), were included as a control to determine the initial inoculum size. The average diameters of each strain were calculated from three independent plates; the results are represented as the mean of diameters ± standard error of the mean (SEM).

## Results and Discussion

### Purification and preparation of the *Td* FlgE G11-M454: 5-FITC ethylenediamine conjugate

*Td* FlgE G11-M454 was employed as the ELISA coat protein and was also used to prepare the FlgE fluorophore protein conjugate that serves as the second protein for the crosslinking reaction. *Td* FlgE G11-M454 lacks part of its D0 domain that facilitates interactions among FlgE subunits; thus, only those proteins that undergo Lal crosslinking remain stably associated^20^. In addition, this variant undergoes crosslinking in solution in the presence of N-ethyl-maleimide (NEM), which aids in converting Cys178 to DHA during the crosslinking reaction^20^. *Td* FlgE G11-M454 is also readily purified recombinantly with high yield, solubility and purity from *E. coli* **(Fig. S1)**, unlike full-length *Td* FlgE which requires refolding from inclusion bodies^20^.

For detection, we chose a fluorophore that could be conjugated to the carboxyl groups of the protein to avoid blocking the ε-amino group of Lys165 involved in the Lal crosslink. 5-FITC ethylenediamine was preferred over 6-aminofluorescein as the former has an extended ethylene linker that would provide greater flexibility for binding to the anti-FITC:HRP conjugate. Anti-FITC:HRP conjugates are commercially available. The conjugation reaction was performed using NHS and EDC in HEPES buffer (pH 7.5), which lacked free amines (see Materials and Methods).

The conjugation of fluorophore 5-FITC ethylenediamine to FlgE was verified through fluorescence detection of the protein conjugate on a plate reader **(Fig. 1B)**. The FlgE:fluorophore conjugate gave an emission maxima at 521 nm (with excitation at 470 nm and cutoff at 495 nm), whereas the non-conjugated protein showed no fluorescence.

**Fig. 1:**
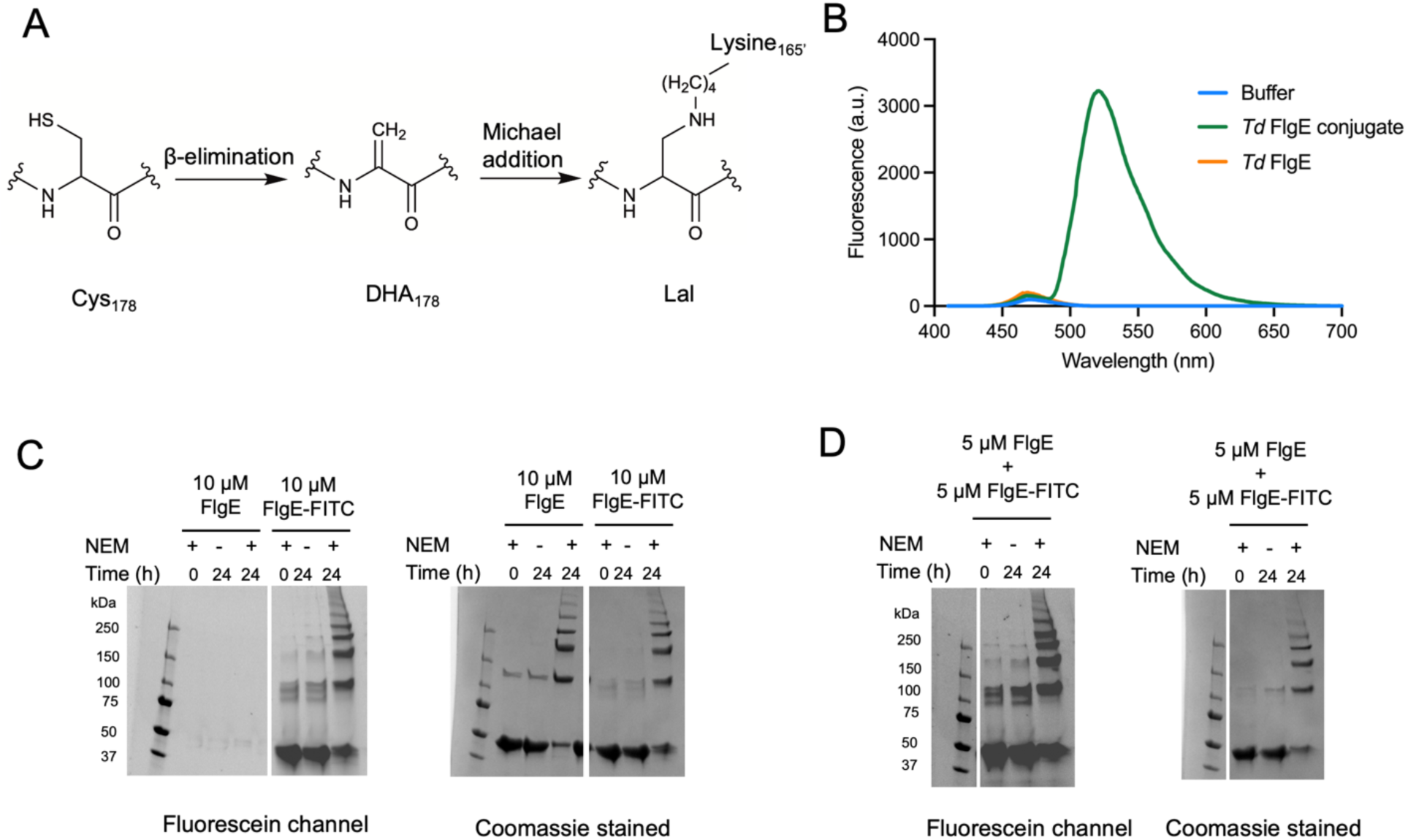
Inhibition of Lal crosslink formation as a basis for an ELISA-type screening procedure. (A) The Lal crosslink forms spontaneously between C178 and K165 through a two-step process involving the intermediate dehydroalanine (DHA). (B) Fluorescence emission spectra of FlgE conjugated to FITC ethylenediamine and non-conjugated protein. 20 μM protein was excited at 470 nm with cut-off at 495 nm and spectra were collected over 410-700 nm. The *Td* FlgE G11- M454:5-FITC ethylenediamine conjugate has an emission maximum at 521 nm. (C) SDS-PAGE gels showing HMWC formation due to Lal crosslinking of FlgE conjugate and non-conjugate imaged both on a ChemiDoc fluorescein channel (left) and after Coomassie-staining (right). Both forms of the protein produce HMWCs, thus indicating that conjugation to 5-FITC ethylenediamine does not affect crosslink formation but does allow for fluorescence detection. (D) SDS-PAGE gels showing HMWC formation of an equimolar mixture of 5 μM conjugated + 5 μM non-conjugated protein as visualized on a ChemiDoc fluorescein channel (left) and after Coomassie-staining (right).

An SDS-PAGE crosslinking assay verified that the FlgE:5-FITC ethylenediamine conjugate could undergo Lal crosslinking. Samples were prepared by incubating 10 μM FlgE or the FlgE:fluorophore conjugate in crosslinking buffer either in the presence or absence of 1 mM NEM for 24 h and then quenching with Laemmli buffer. After running the samples on a Tris-glycine SDS-PAGE gel and staining with Coomassie, both non-conjugated and conjugated FlgE showed comparable amounts of HMWCs, thus confirming that fluorophore conjugation did not affect Lal formation. Prior to staining with Coomassie, the same gel was also imaged using the fluorescein application (excitation source: Blue Epi illumination, emission filter: 530/28) on a Bio-Rad ChemiDoc MP imager. When imaged under this channel, the same HMWC protein bands seen after Coomassie-staining, are observed only for the FlgE:fluorophore conjugate **(Fig. 1C)**. An equimolar protein mixture of 5 μM fluorophore-conjugated + 5 μM non-conjugated protein shows a similar pattern and intensity of HMWC bands as the gels in Fig. 1C **(Fig. 1D)**. This similarity in signal indicates that the FlgE:fluorophore conjugated protein and the non-conjugated protein are capable of crosslinking to each other, as if they did not, the number and intensity of bands would be lower than that in Fig. 1C.

SDS-PAGE gel assays were also performed with an equimolar mixture of 5 μM of C178A FlgE:fluorophore conjugated protein + 5 μM of WT non-conjugated protein **(Fig. S2)**. The lower crosslinking yields of the mixed sample compared to WT alone reflects that the 178 position in the C178A variant cannot undergo the Lal reaction, and only the Lys in this variant can react. Nevertheless, the two populations of proteins can crosslink with each other, as HMWCs can be seen in both the fluorescein channel and Coomassie-stained gel to the same extent as with equimolar mixtures of non-conjugated WT and C178A proteins.

### Development of the ELISA for HTS

The optimized ELISA was performed by coating Nunc MaxiSorp strips/plates with *Td* FlgE G11- M454 followed by blocking with 2% non-fat dry milk (NFDM). *Td* FlgE G11-M454:5-FITC ethylenediamine conjugate was added to the wells in crosslinking buffer followed by NEM. The crosslinking reaction was allowed to proceed for 24 h. FlgE crosslinking was followed by incubation with anti-FITC:HRP antibody for 1 h. Color development was initiated by adding ABTS solution and the reaction was quenched after 15 min with 1% SDS **(Fig. 2)**.

**Fig. 2:**
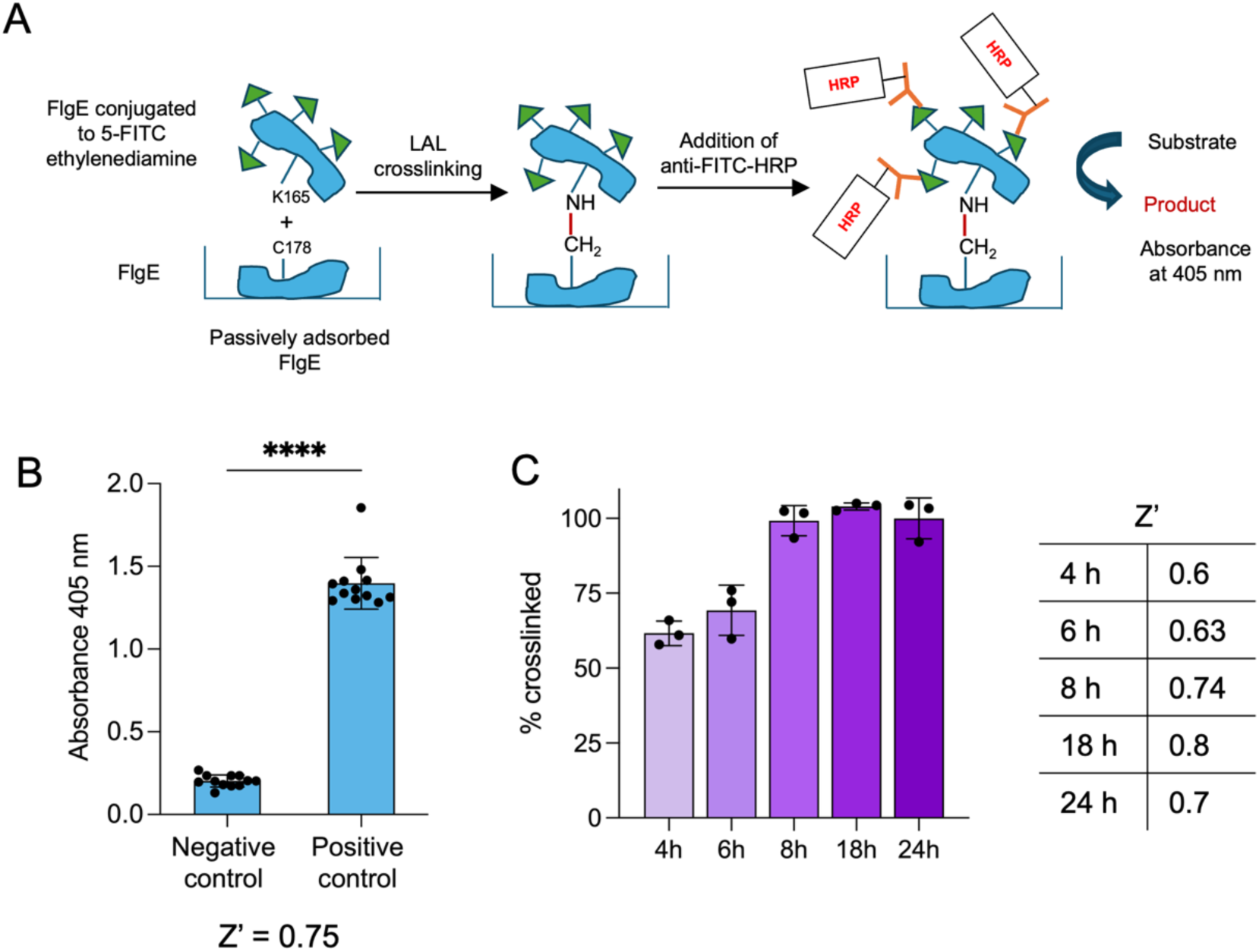
ELISA for identification of Lal crosslinking inhibitors (A) Schematic of the ELISA. *Td* FlgE G11-M454 was coated on Nunc MaxiSorp strips through passive adsorption and strips were subsequently blocked in 2% NFDM. *Td* FlgE G11-M454: 5- FITC-ethlynediamine conjugate was added to the wells in crosslinking buffer with 1 mM NEM. Lal crosslinking was allowed to proceed for 24 h. Strips were incubated with anti-FITC-HRP for one hour, washed, and then ABTS solution containing H2O2 was added. Color development occurred in proportion to the amount of crosslinking. (B) Signal generated by the positive and negative control. The negative control involved not coating the strips with FlgE, but proceeding with all subsequent steps of blocking, i.e. adding conjugate, incubating with antibody and developing color. The positive control involved following the optimized conditions in (A), including the first step of coating the strips with FlgE, followed by all the steps listed above. Consistent Z’ values of > 0.65 were obtained when using freshly coated strips. Results are shown in absorbance units (a.u.) as the mean value ± standard deviation (S.D.) on n = 12 technical replicates. Statistical significance was calculated using a two-tailed t-test (*p* < 0.05*, 0.01** and 0.001***). (C) Time course of crosslink formation with corresponding Z’ values. The conjugate was added to coated strips and the crosslink reaction was allowed for proceed for 4, 6, 8, 18 and 24 h. Near complete signal is obtained after 8 h. Results were shown as the mean value ± standard deviation (S.D.) on n = 3 technical replicates.

During the development of the ELISA, the type of blocking solution, temperature of the crosslinking reaction, antibody dilutions, and time of color development were varied and optimized. We aimed to generate an OD of 1.2-1.5 in 15 min of color development prior to stopping the reaction. Anti-FITC:HRP conjugate was tested at 10k, 20k 40k, and 80k dilutions of the stock (Invitrogen, RRID identifier AB_930865), and the one which gave an OD of 1.2-1.5 for the 15 min color development was selected as an initial dilution **(Fig. S3)**.

Similarly, four different blocking solutions, 1% BSA, 2% non-fat dry milk (NFDM) in PBS, protein-free (TBS) blocking buffer (ThermoFisher), and 5% calf serum (Gibco Laboratories) were evaluated to determine the optimal blocking solution with assay conditions of 1:20000 antibody dilution and 15 min of color development **(Fig. S4)**. The protein-free blocking buffer and 1% BSA gave higher background in non-crosslinking buffer conditions and were eliminated from further experimentation. Blocking solutions containing NFDM and calf serum gave a similar low background. With these considerations and cost assessment, 2% NFDM was used as a blocking buffer throughout the assay development process. The temperature and antibody dilution were then further optimized by performing the crosslinking reaction at 4°C, 23°C (RT) and 37°C, with two antibody dilutions of 1:7500 and 1:15000 **(Fig. S5)**. The reaction temperature of 23°C and antibody dilution of 1:15000 gave the highest fold-change between negative and positive controls and generated an OD of ∼1.4 after 15 min of color development. The data also indicate that increasing the anti-FITC:HRP concentrations (lower dilutions) generates a much higher signal readout (50- 90% higher).

A negative control, wherein the wells were not coated with FlgE but all other assay steps were performed, accompanied all test conditions. This negative control ensured that the FlgE:fluorophore does not form crosslinks with Lys or Cys residues in the blocking protein, even under crosslinking conditions (buffer composition of 40 mM Tris pH 8.5, 160 mM NaCl, 1 M ammonium sulfate, 1 mM NEM). An absolute blank without coat protein or FlgE conjugate, i.e., with wells blocked, antibody added, and color development showed near zero values, verifying that the blocking solution works well and that the anti-FITC:HRP antibody does not interact with the blocking protein. An additional control in which the FlgE:fluorophore was added to coated and blocked plates under buffer conditions not conducive to crosslinking (in buffer lacking ammonium sulfate and NEM i.e. 40 mM Tris pH 8.5, 160 mM NaCl) generated low signal as well, further validating the specificity of the ELISA **(Fig. S6)**.

The positive crosslinking control was executed by coating the wells with FlgE, blocking the wells, initiating the crosslinking reaction by adding the FlgE:fluorophore conjugate in crosslinking buffer followed by the addition of NEM, and then adding the detection antibody and developing color. The positive crosslinking control (with FlgE coat) produced a 5-7-fold increase over the non-coated (negative crosslinking) control **(Fig. 2B)**. Z’ values of 0.55 to 0.75 were routinely obtained for the final optimized assay, with coated and blocked strips used within 7-10 days giving higher Z’ values (> 0.70) than the strips that were stored for longer periods of time (0.6).

### Time course of Lal crosslink formation using the ELISA

*Td* FlgE Lal crosslinking routinely takes 24-48 h to achieve maximum crosslinking in various assays^19,20,34^. Such long incubation times are a drawback in HTS; for example, they exacerbate plate edge effects. Unlike the NanoLuc assay, the ELISA uses an enzyme-mediated signal amplification step that potentially increases sensitivity at shorter incubation times. To test this possibility, the Lal crosslinking reaction was allowed to proceed at various time intervals, and the signals generated were compared to those obtained after 24 h of crosslinking **(Fig. 2C)**. Crosslinking assays performed at 4, 6, 8, 18 and 24 h under the optimized assay conditions showed that a near maximum signal is obtained within 8 h, thus indicating that the assay time could be reduced substantially from previous methods. Because of the signal amplification system in the ELISA, measurable Lal formation was apparent even as crosslinking times were lowered to about six hours (60-70% crosslinking), which rivals the earliest formation of faint HMWC bands in the electrophoretic assay^4^. Thus, the ELISA can lower assay times for HTS from 24-28 h to 8-10 h.

### Effects of hexachlorophene (HC) and triclosan (TC) on Lal crosslinking

We previously identified two compounds, hexachlorophene (HC) and triclosan (TC), which inhibited Lal crosslinking when a small library of 727 compounds was screened using the NanoLuc assay^5^. Both compounds also showed inhibition in the *in vitro* SDS-PAGE crosslinking assay with *Td* FlgE and were active to varying degrees for FlgE from other spirochete species. HC was found to be a more potent inhibitor with an IC50 of 9 μM compared to 48 μM for TC. HC was also found to block FlgE crosslinking *in vivo* and affect *T. denticola* motility^5^.

In the ELISA assay, TC was found to be a much weaker inhibitor of HMWC formation than HC **(Fig. 3A)**. Only about 60% inhibition was observed at concentrations as high as 1 mM. TC was thus excluded from further studies. Similar to the NanoLuc and SDS-PAGE assays, HC acted as a stronger inhibitor than TC with an IC50 value of ∼40-50 μM, which is higher than that found by the *in vitro* SDS-PAGE assay (∼8-10 μM). This difference may reflect the very different concentrations of FlgE being used in the two assays (10-100x higher concentrations in the SDS-PAGE assays) and the fact that one of the crosslinking partners is immobilized in the ELISA.

**Fig. 3:**
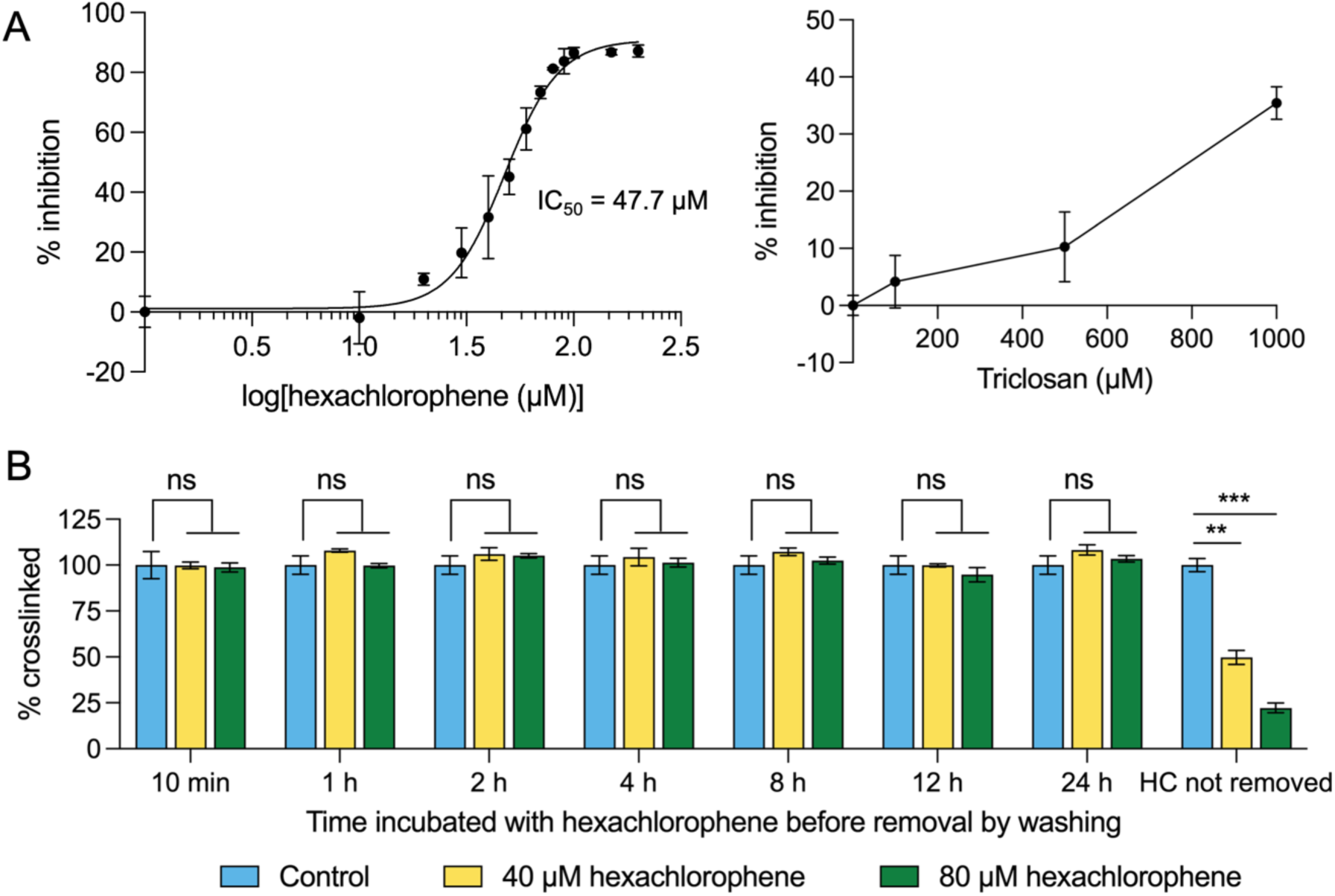
Inhibition of crosslinking by hexachlorophene and triclosan. (A) Dose-response curve generated by the ELISA with 0-200 μM HC added during the Lal crosslinking step (left). An IC50 of 47.7 μM was obtained for hexachlorophene. Triclosan is a weaker inhibitor of crosslinking showing only ∼35% inhibition at 1000 μM (right). Data was plotted from 3 technical replicates. (B) Exchanging HC in wash steps abrogates inhibition. Coated strips were incubated with HC (40 μM and 80 μM) for 0.17 - 24 h before washing strips to remove unbound HC and then proceeding with the crosslinking reaction by adding FlgE conjugate. Strips for which HC was not removed and was present for the entire crosslinking reaction along with conjugate were used as a comparison. Results were shown as the mean value ± standard deviation (S.D.) on n = 3 technical replicates. Statistical significance was calculated using a one-way ANOVA with post-hoc Dunnett’s test (*p >*0.05 ns*, p* < 0.05*, 0.01** and 0.001***).

The NanoLuc assay indicated that HC inhibition was to a degree irreversible, consistent with a covalent interaction of HC with the protein, perhaps through the formation of a DHA-HC adduct. We thus evaluated the mode of action of HC on Lal formation in the ELISA assay, which unlike the NanoLuc, does not involve the pre-conversion of Cys178 to DHA, which is accompanied by a substantial change in protein conformation^20^. In the ELISA format, we included an additional step in which the FlgE-coated strips were first treated with 200 μM HC for different time intervals (10 min and 1-24 h) **(Fig. 3B)**. After washing the strips, FlgE:fluorophore conjugate was added, and the assay was performed according to the standard protocol. Pre-incubating the strips with HC prior to its removal did not affect the formation of HMWCs. In contrast, strips similarly pretreated with HC, its subsequent removal by washing, and then re-addition to 40 and 80 μM gave a similar inhibition pattern as seen with the standard protocol. In addition, pre-incubation of HC and the FlgE coat protein with and without NEM under different buffer conditions prior to coating the plates did not alter the inhibition behavior **(Fig. S8)**.

Thus, in this format, it seems unlikely that HC covalently modifies Cys178 prior to Lal formation. In principle, crosslinking can occur in either direction, with the participating DHA on either the adsorbed protein or the FITC-conjugated protein. Furthermore, Lal crosslinking also depends on Asn179 and proceeds through β-elimination of Cys178 to form sulfide and the intermediate DHA^20^. Thus, it may be that the DHA-HC adduct forms only after the Asn179 from the FlgE:fluorophore conjugate in complex with the adsorbed protein facilitates the conversion of Cys178 to DHA. The availability of DHA in this reaction may be quite transient, unlike in the NanoLuc assay, where it is pre-formed. Alternatively, the NEM-mediated conversion of Cys178 to DHA, which is not a feature of the NanoLuc assay, may interfere with covalent modification by HC. That said, we attempted to isolate a tryptic peptide of FlgE containing an HC, or HC-derived adduct to no avail and thus the irreversible nature of the interaction in the NanoLuc may result instead from a slow off rate of the compound bound to the DHA form.

### Dopamine (DOPA) as a crosslinking activator

Dopamine (DOPA) and several dopamine-related catecholamines appear to increase Lal crosslinking. To test the effect of DOPA on the ELISA, the assay was run using the optimized protocol with different concentrations of DOPA in the absence and presence of 80 μM HC **(Fig. S9)**. At lower concentrations (1-4 μM), DOPA acted as an activator, increasing the Lal formation by approximately 5-14%. However, at higher concentrations (> 30 μM), DOPA lowered Lal formation by almost 20%. This is in contrast to the behavior observed with NanoLuc assay where DOPA increased Lal formation in a dose-dependent manner. However, the DOPA effect in the NanoLuc assay was attributed to DOPA both accelerating crosslinking and also causing non-specific protein aggregation^5^. Enhanced aggregation likely facilitates association of the split luciferase reporter, increasing signal. In contrast, protein aggregation would sequester the soluble FlgE:fluorophore conjugate in the ELISA, causing less protein to be available to crosslink to the adsorbed FlgE, and be removed during the wash step, lowering signal. This is likely to be the reason that higher concentrations of DOPA showed inhibitory effects on Lal formation in the ELISA, but not the NanoLuc assay.

The non-aggregate associated DOPA effect that increased both the NanoLuc and the ELISA signal may be related to the ability of DOPA to form cysteinyl adducts^35^, which would facilitate conversion to DHA, much like NEM. Indeed, DOPA partially reversed HC inhibition of Lal crosslinking. The inhibitory effects of HC on Lal crosslinking in the presence of 100 μM DOPA were lowered by almost 35-40% **(Fig. S9)**. An even more striking effect of DOPA is observed in the absence of NEM, where the addition of 10 μM DOPA to conjugate with 80 μM HC led to a 75% increase in crosslinking over conjugate with 80 μM HC alone.

### Screening of NIHCC compounds

The optimized ELISA assay was used to screen the NIHCC library. Compounds at a final assay concentration of 200 μM were added with the FlgE:fluorophore conjugate in crosslinking buffer to coated and blocked strips. NEM was added after 30 min of incubation with the compounds. After assay assessment, compounds that showed inhibition ≥45% compared to the control were validated using the SDS-PAGE gel-based assay **(Fig. 4A,B)**. Densitometric analysis was performed using ImageJ on the HMWC bands obtained on the gels to roughly quantify the amount of inhibition **(Fig. 4C)**. Based on the ELISA and densitometric analysis, three compounds were identified as effective inhibitors: hexachlorophene, honokiol and zafirlukast **(Fig. 4D)**. HC gave the highest inhibition with 82.3% in the ELISA and 78.4% in the gel compared to control. Honokiol and zafirlukast inhibited Lal to a lesser degree but still show reduction of the trimer, tetramer and higher order oligomers in the gel.

**Fig. 4:**
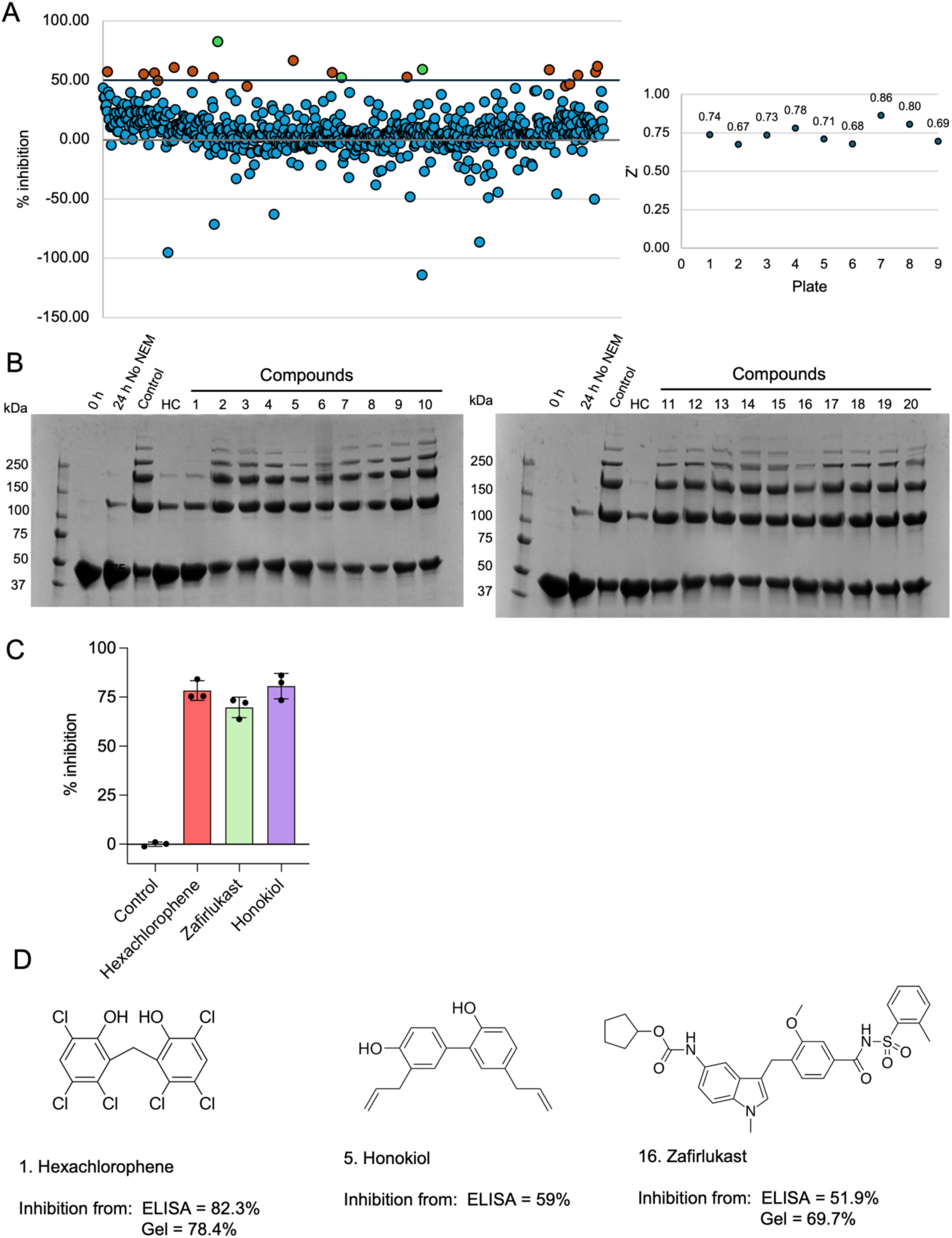
ELISA screening of the NIHCC compound library (A) Scatter plot showing crosslink inhibition by each compound of the collection compared to no-compound control (left). Dots in red show compounds causing ≥ 45 % inhibition. These compounds were further studied using the SDS-PAGE gel assay. The three compounds hexachlorophene, zafirlukast and honokiol that were selected based on the ELISA and initial confirmatory SDS-PAGE assay are designated by green dots. Several compounds were also found to increase crosslinking as shown by the negative % inhibition values. Z’ values obtained for each plate are shown (right). (B) SDS-PAGE assay of the 20 compounds showing ≥ 45 % inhibition in the ELISA. (C) Densitometric analysis by ImageJ on the best hits from the SDS-PAGE assay. The intensities of all HMWC bands on the gel were combined and compared to those of the control. Honokiol showed artificially high inhibition because of protein precipitation that limited the amount of protein entering the gel lanes. (D) Structures of compounds 1, 5 and 16 from the SDS-PAGE assay and the % inhibition obtained from the ELISA and gel-based assays.

Honokiol has some structural similarity with hexachlorophene, both being biphenols. HC, however, is an antiseptic and has relatively high toxicity (LD50 of 55-65 mg/kg in rats)^36^. Unlike HC, both honokiol and zafirlukast are non-toxic and thus are likely to be more amenable to cellular studies^37–41^.

Several compounds identified by the ELISA were also found to activate crosslinking **(Fig. S10)**. These compounds could be separated into groups based on structural similarity. Surprisingly, many of these activators were found to belong to different antibiotic families. Two cephalosporins caused the highest increase in crosslinking signals with values of ∼200% of the no-compound control. Several tetracyclines and plant polyphenols were also identified to increase crosslinking.

It is also interesting to note that rifampicin was one of the antibiotic activators identified, as spirochetes are naturally resistant to it^42,43^.

### Inhibition of crosslinking by zafirlukast and honokiol

Zafirlukast and honokiol were the most promising new hits obtained after both the ELISA HTS and the secondary SDS-PAGE assays. To determine IC50 values, the ELISA assay was performed on a dilution series of 0-300 μM of zafirlukast and 0-750 μM of honokiol **(Fig. 5A)**. An IC50 of 92 μM was obtained for zafirlukast, indicating that this compound may be used as a starting point for structure-activity modifications to improve inhibition. Because percent of inhibition with honokiol plateaued at ∼75%, the data was normalized to this value to calculate an IC50 of 118 μM. The plateau effect likely derives from the low solubility of honokiol and its tendency to cause protein precipitation. SDS-PAGE gels of Lal inhibition were also performed with increasing concentrations of these compounds **(Fig. 5B)**. To evaluate if the ELISA could identify a compound with activity against *T. denticola* cells, we tested the most potent new inhibitor, zafirlukast for its ability to block FlgE crosslinking of growing *T. denticola* cells and impair their motility in swim plates. Like hexachlorophene, we found that zafirlukast also increases low molecular weight FlgE species by Western blot, and correspondingly reduces motility, at concentrations that do not affect cell growth (**Fig. S11**).

**Fig. 5:**
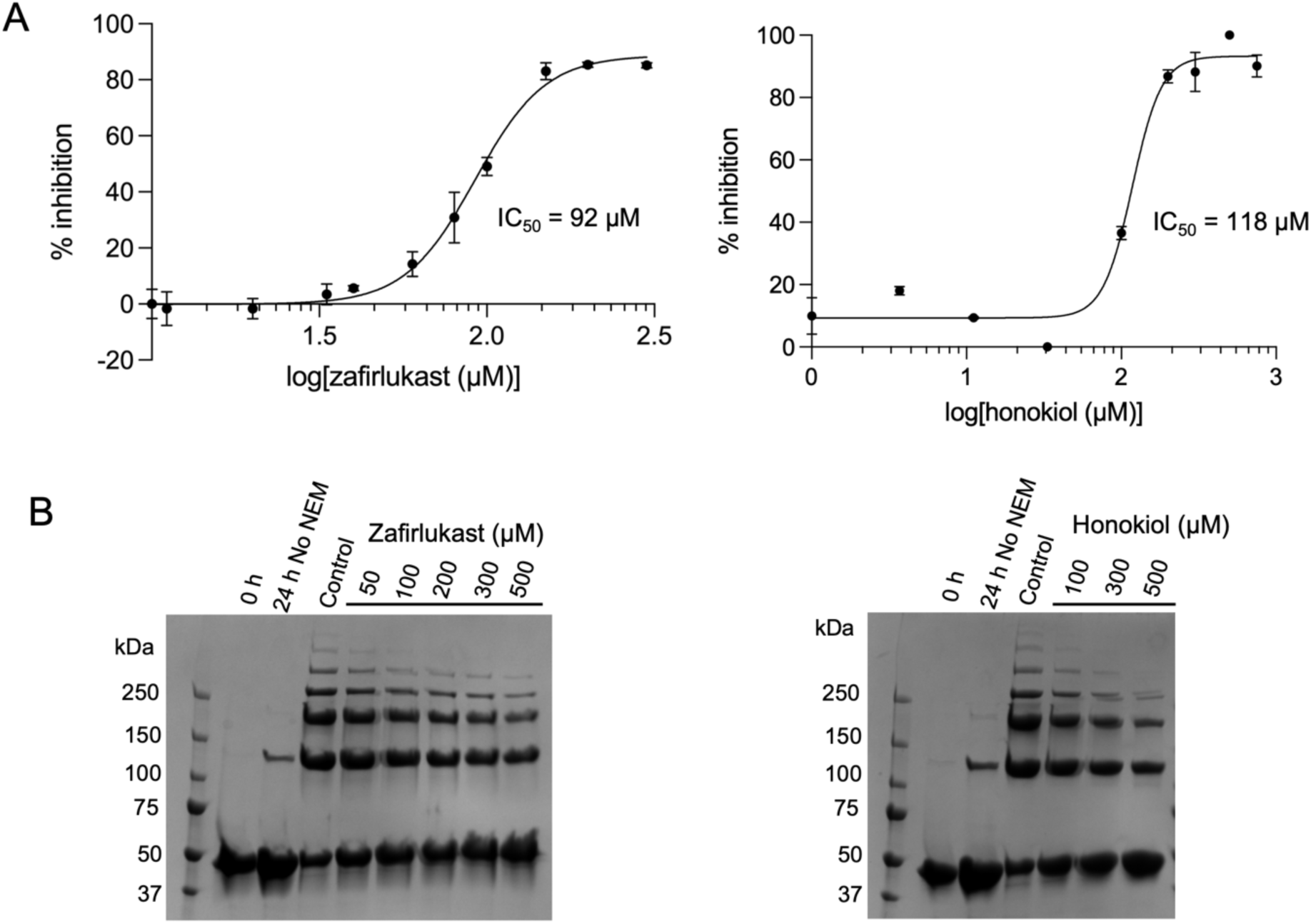
Inhibition of crosslinking by zafirlukast and honokiol. (A) Dose-response curve generated by the ELISA with 0-300 μM of zafirlukast (left) and 0-750 μM of honokiol (right) added during the Lal crosslinking step. An IC50 of 92 μM was obtained for zafirlukast and 118 μM for honokiol on fitting the results to a four-parameter equation (see Materials and Methods). The data for honokiol was normalized as inhibition plateaued at ∼75%. Data was plotted from 3 technical replicates. (B) SDS-PAGE assay showing the inhibition of Lal crosslinking with increasing concentration of inhibitor. There is a decrease in the intensities of HMWCs (> 250 kDa) on treatment with the compounds.

## Conclusions

Lysinoalanine (Lal) crosslinking between Cys178 and Lys165 of FlgE subunits via the dehydroalanine (DHA) adduct is essential for motility and thereby virulence of spirochetes. Lal thus offers a valid target for high throughput screening of large compound libraries to develop potential new drug leads to treat spirochete infections. We here report the development and optimization of a heterogeneous sandwich ELISA assay that employs a tethered wild-type FlgE protein to closely mimic conditions during hook biogenesis. By lowering the crosslinking reaction to 8 h, which allows for full Lal conversion (Fig. 2), we could complete the ELISA assay in under 10 hours. As the ELISA is a signal amplification enzyme assay, one could further modify assay conditions such as increasing the concentration of the FlgE:5-FITC ethylene diamine conjugate as well as the anti-FITC:HRP conjugate, decreasing incubation time with the anti-FITC:HRP conjugate to 30 min or less, and increasing the color development time from 15 min to 30 min. A more rapid assay would be beneficial by mitigating issues such as plate edge effects, protein and reagent instability and environmental variation.

Hexachlorophene (HC) was verified to be a potent inhibitor (IC50 ∼50 μM) of the Lal crosslinking reaction. Dopamine (DOPA) acted as an activator of crosslinking reaction at lower concentrations (1-4 μM) and also reversed the HC inhibition of Lal crosslinking. Screening of a small library of clinically approved NIH compounds with the ELISA identified two new compounds, zafirlukast and honokiol as inhibitors, the former of which shows activity against *T. denticola* cells. The assay also detected several activator molecules, which surprisingly belong to some well-known commonly used families of antibiotics. In addition to advancing potential therapeutics against spirochetes, this study demonstrates the utility of an ELISA-based sandwich assay for high-throughput monitoring of protein-protein interactions, a desirable, but often difficult process to target. In this case, FlgE benefits from the formation of a covalent crosslink between the immobilized and fluid phase proteins; however, protein-protein interactions with KD < 1 μM (off rates < 0.1 sec^-1^) would also be applicable to this same approach.

## Supporting information

none

## Acknowledgements

This work was supported by NIAID grants R01AI148844 to BRC, R01AI078958 to CL and the Chemical Biology Interface NIH training grant T32GM138826 to MD.

## Data Availability Statement

The data that support the findings of this study are available in the Supporting Information.

