## Supplementary material for "An ELISA for discovering protein-protein interaction inhibitors: blocking lysinoalanine crosslinking between subunits of the spirochete flagellar hook as a test case": none

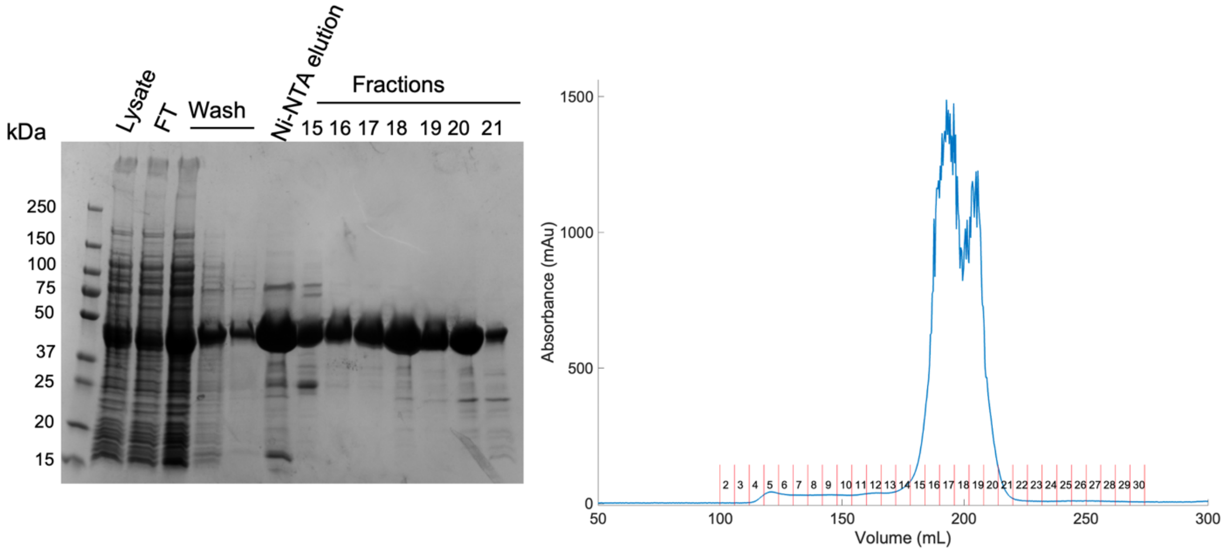

Fig. S1: Purification of recombinant *Td* FlgE G11-M454 from *E. coli*

Coomassie-stained SDS PAGE gel (left) and FPLC SEC chromatography trace (right) for the purification of *Td* FlgE G11-M454 from 4 L of *E. coli* cell culture growth. Fractions 18-20 were pooled and used for the WT FlgE ELISA coat protein and to make the FlgE:5-FITC ethylenediamine conjugate.

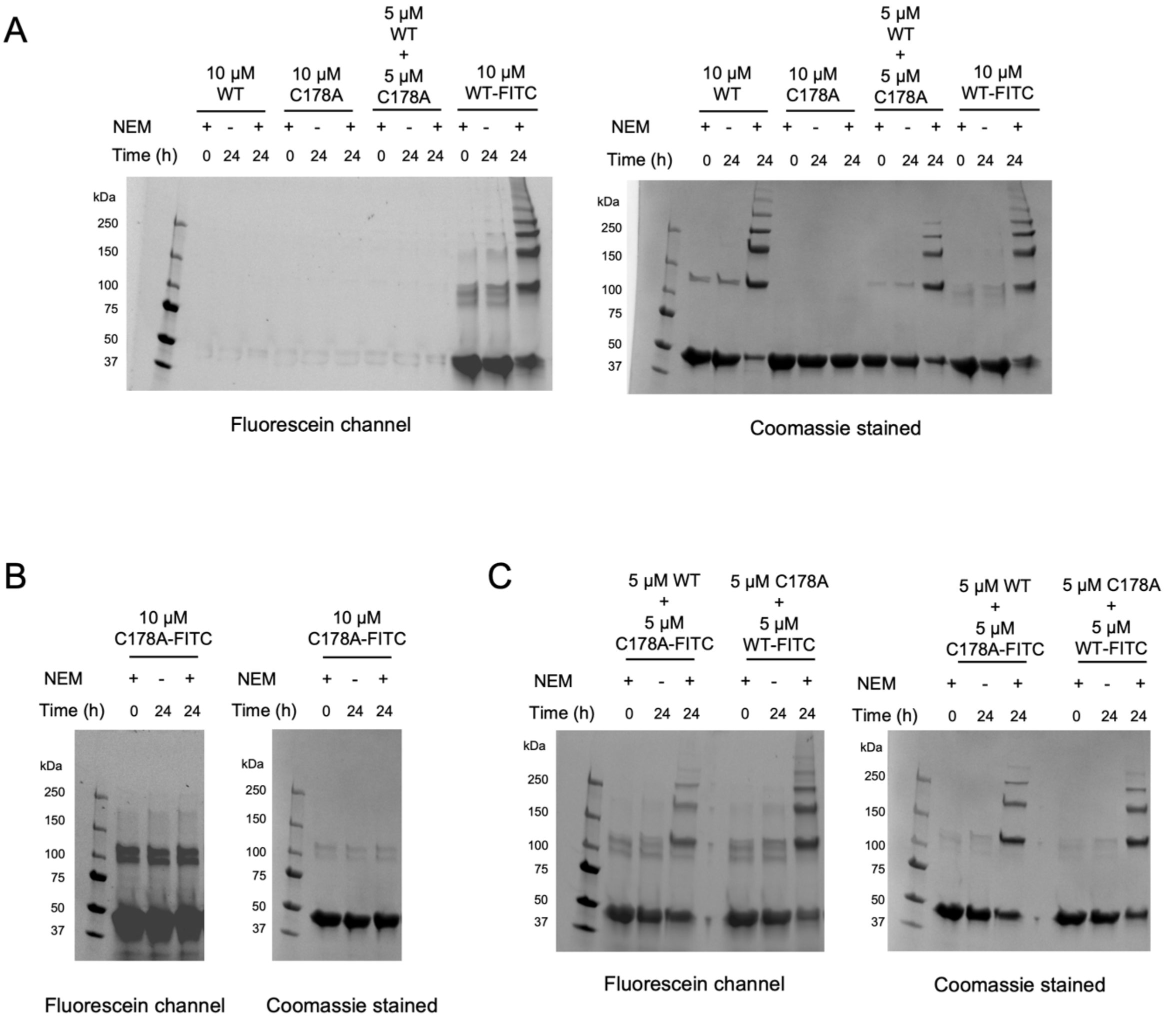

Fig S2: SDS-PAGE gels on Lal crosslink formation in 5-FITC ethylenediamine conjugated and non-conjugated WT *Td* FlgE G11-M454 and C178A mutant.

FlgE:5-FITC ethylenediamine conjugation has been represented as WT-FITC or C178A-FITC on the gel for ease of labeling.

(A) 10 μM of WT and WT-FITC protein both show comparative levels of HMWC formation on the gel, indicating that 5-FITC ethylenediamine conjugation does not affect crosslinking. An equimolar mixture of 5 μM of WT + 5 μM of C178A protein shows lesser crosslinking than WT alone, as the Cys in the C178A mutant cannot undergo the Lal reaction, thus decreasing the extent of HMWC formation. In this case, crosslinks can form between two WT proteins, or between Lys in the C178A mutant and Cys in the WT protein.

(B) The C178A-FITC protein cannot crosslink with itself.

(C) An equimolar mixture of 5 μM of WT + 5 μM of C178A-FITC protein confirms that crosslinking can occur between the Lys in C178A-FITC mutant and Cys in the WT protein as HMWCs can be seen in both the fluorescein channel and after staining with Coomassie to same extent as in (A). The same result is observed with reacting the C178A mutant with WT-FITC. Thus, the conjugation of the fluorophore 5-FITC ethylenediamine does not affect FlgE crosslinking.

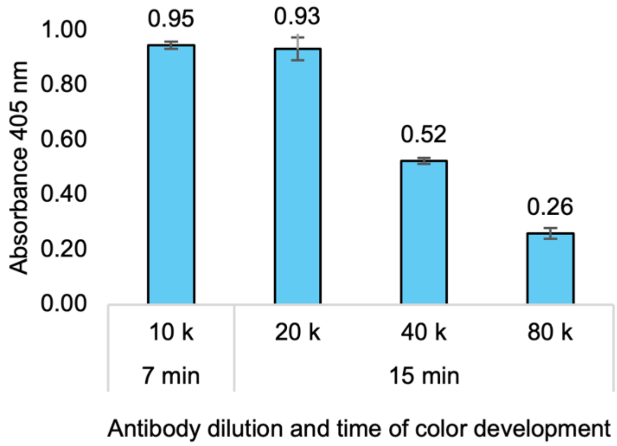

Fig. S3: HRP antibody dilution ratios for optimization of ELISA signals

Antibody dilutions of 1:10,000, 1:20,000, 1:40,000 and 1:80,000 were used and color development was quenched at either 7 min or 15 min before measuring the absorbance. The ratio of absorbance obtained at each antibody dilution can be correlated to the amount of antibody added (e.g. absorbance of 0.26 at 80,000x antibody dilution is half that obtained at 40,000x dilution), thus indicating that the ELISA can be used quantitatively to measure the degree of conjugated FlgE obtained from crosslinking.

For this optimization experiment, the plate was coated with FlgE (100 µg/ml in PBS, pH 7.2; 100 µL/well) for 24 h at 4°C and blocked with 1% BSA for 4 h at room temperature. FlgE:fluorophore conjugate (10 µg/mL, 100 µL/well) in crosslinking buffer (40 mM Tris pH 8.5, 160 mM NaCl, 1 M ammonium sulfate) was added to the wells along with 1 mM NEM. The Lal crosslinking reaction was performed for 18 h at room temperature, followed by incubating with anti-FITC HRP conjugated antibody at the dilutions listed above, for 30 min prior to color development. ABTS substrate was added for 7 min or 15 min and subsequently quenched with 1% SDS prior to reading absorbance at 405 nm.

Results are shown in absorbance units (a.u.) as the mean value ± standard deviation (S.D.) on 3 technical replicates for each antibody dilution.

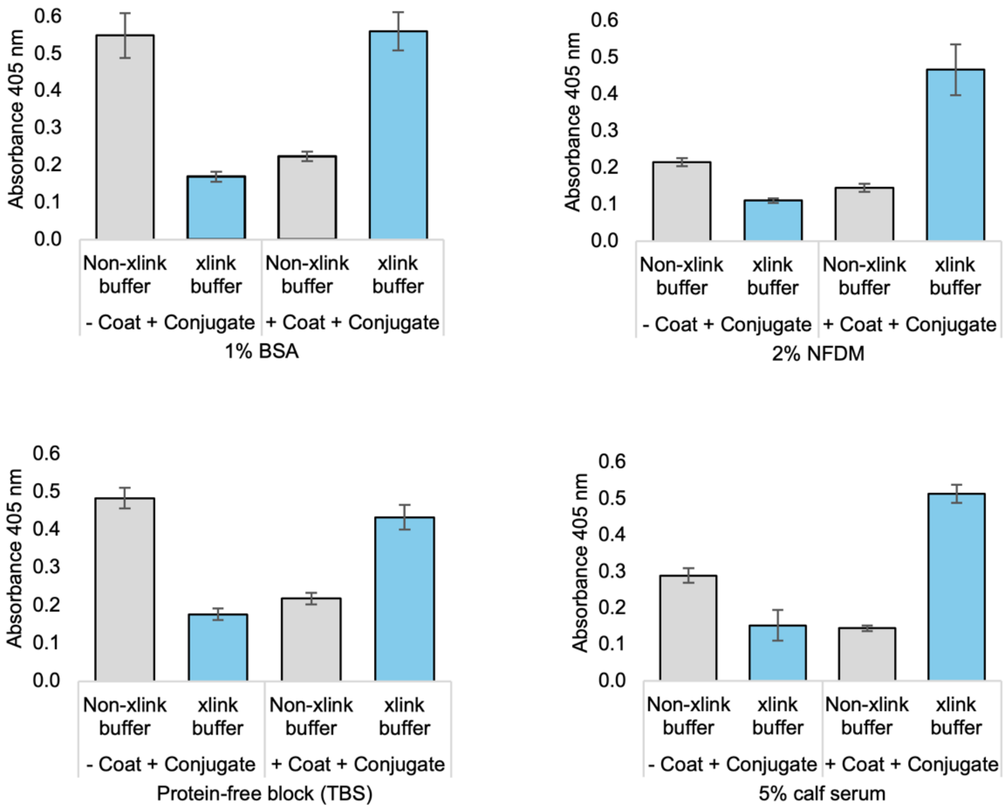

Fig. S4: Optimization of blocking buffer using an antibody dilution of 1:20000

Four blocking buffers were studied – 1% BSA, 2% NFDM, 5% calf serum and Pierce™ protein-free blocking buffer (TBS) under the negative and positive control conditions. BSA and protein-free block gave higher signals in non-coated plated under non-crosslink (non-xlink) buffer conditions lacking ammonium sulfate (40 mM Tris pH 8.5, 160 mM NaCl) and thus were eliminated from the assay.

For this optimization experiment, the plate was coated with FlgE (100 μg/ml in PBS, pH 7.2; 100 μL/well) for 24 h at 4°C and blocked with the different blocking solutions listed above for 4 h at room temperature. FlgE:fluorophore conjugate (10 μg/mL, 100 μL/well) in either crosslink (xlink) buffer (40 mM Tris pH 8.5, 160 mM NaCl, 1 M ammonium sulfate) or non-crosslink (non-xlink) buffer (40 mM Tris pH 8.5, 160 mM NaCl) was added to the wells along with 1 mM NEM. The Lal crosslinking reaction was performed for 18 h at 4°C, followed by incubating with anti-FITC HRP conjugated antibody, for 1 h prior to color development. ABTS substrate was added for 15 min and subsequently quenched with 1% SDS prior to reading absorbance at 405 nm.

Results are shown in absorbance units (a.u.) as the mean value ± standard deviation (S.D.) on 6 technical replicates.

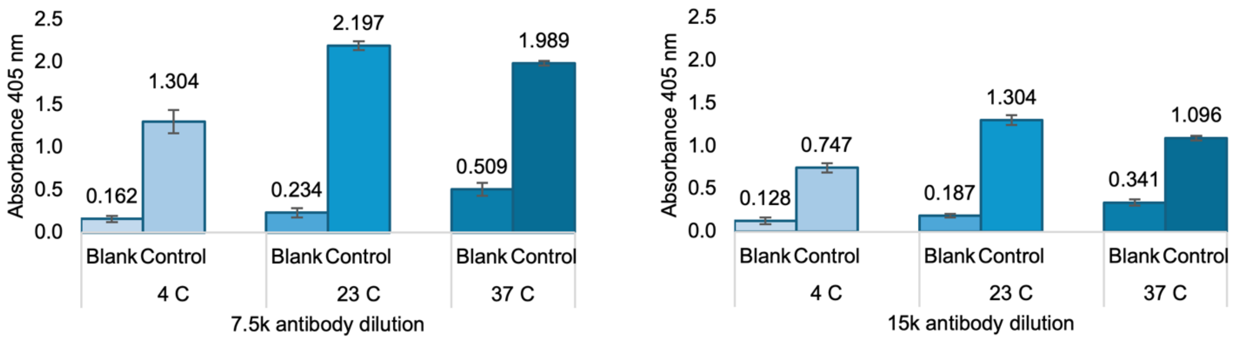

Fig. S5: Optimization of the temperature of the crosslinking reaction and antibody dilution

The crosslinking reaction was allowed to proceed for 24 h at 4, 23, or 37°C. Two different antibody dilutions of 1:7500 and 1:15000 were used prior to the color development step. We aimed to achieve an absorbance of ~1.5 after ~15 min of color development. The conditions chosen for HTS involved performing the crosslinking reaction at room temperature and using an antibody dilution of 1:15000.

For this optimization experiment, the plate was coated with FlgE (100 µg/ml in PBS, pH 7.2; 100 µL/well) for 24 h at 4°C and blocked with 2% non-fat dry milk for 4 h at room temperature. FlgE:fluorophore conjugate (10 µg/mL, 100 µL/well) in crosslinking buffer (40 mM Tris pH 8.5, 160 mM NaCl, 1 M ammonium sulfate) was added to the wells along with 1 mM NEM. The Lal crosslinking reaction was performed for 24 h at 4°C, 23°C or 37°C followed by incubating with anti-FITC HRP conjugated antibody at either 1:7500 or 1:15000 dilution for 1 h prior to color development. ABTS substrate was added for 15 min and subsequently quenched with 1% SDS prior to reading absorbance at 405 nm.

Results are shown in absorbance units (a.u.) as the mean value ± standard deviation (S.D.) on 4 technical replicates.

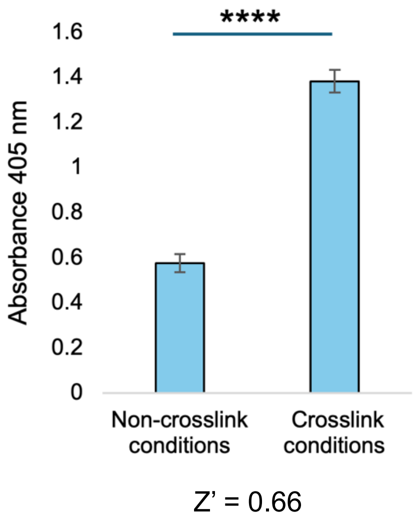

Fig. S6: ELISA performed under crosslinking and non-crosslinking conditions

Signal generated by the ELISA with the Lal crosslink reaction allowed to proceed in non-crosslinking conditions (40 mM Tris pH 8.5, 160 mM NaCl) and crosslinking conditions (40 mM Tris pH 8.5, 160 mM NaCl, 1 M ammonium sulfate, 1 mM NEM) was plotted.

To obtain high levels of crosslinking in solution, as evidenced by the SDS-PAGE gel assay, both ammonium sulfate and NEM are required. The ELISA was performed by adding the FlgE:5-FITC fluorophore conjugate either in buffer conducive to crosslinking (40 mM Tris pH 8.5, 160 mM NaCl, 1 M ammonium sulfate, 1 mM NEM) or in buffer not conducive to crosslinking (40 mM Tris pH 8.5, 160 mM NaCl).

The following assay conditions were used in this experiment: The plate was coated with FlgE (100 µg/ml in PBS, pH 7.2; 100 µL/well) for 24 h at 4°C and blocked with 2% non-fat dry milk for 4 h at room temperature. FlgE:fluorophore conjugate (10 µg/mL, 100 µL/well) was added to the wells either in crosslink conditions (40 mM Tris pH 8.5, 160 mM NaCl, 1 M ammonium sulfate, 1 mM NEM) or non-crosslink conditions (40 mM Tris pH 8.5, 160 mM NaCl). The Lal crosslinking reaction was performed for 24 h at room temperature followed by incubating with anti-FITC HRP conjugated antibody at 1:15000 dilution for 1 h prior to color development. ABTS substrate was added for 15 min and subsequently quenched with 1% SDS prior to reading absorbance at 405 nm.

Results are shown in absorbance units (a.u.) as the mean value ± standard deviation (S.D.) on n = 3 technical replicates. Statistical significance was calculated using a two-tailed t-test ($p < 0.05$*, 0.01** and 0.001***).

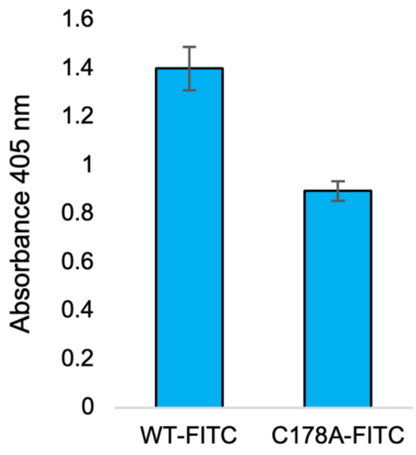

Fig S7: ELISA signal generated by using WT FlgE:5-FITC ethylenediamine conjugate (labeled as WT-FITC) and FlgE C178A:5-FITC ethylenediamine conjugate (labeled as C178A-FITC) on plates coated with WT FlgE

Except for the nature of the FlgE:fluorophore conjugate, the two assays were performed under the same conditions. There is an overall a reduction in the signal obtained with C178A-FITC instead of WT-FITC as only the Lys residue in the C178A variant can crosslink with WT FlgE coated on the plate, and higher-order oligomers cannot form between the WT and C178A proteins.

The following assay conditions were used in this experiment: The plate was coated with FlgE (100 µg/ml in PBS, pH 7.2; 100 µL/well) for 24 h at 4°C and blocked with 2% non-fat dry milk for 4 h at room temperature. Either WT or C178A FlgE:fluorophore conjugate (10 µg/mL, 100 µL/well) in crosslinking buffer (40 mM Tris pH 8.5, 160 mM NaCl, 1 M ammonium sulfate) was added to the wells along with 1 mM NEM. The Lal crosslinking reaction was performed for 24 h at room temperature followed by incubating with anti-FITC HRP conjugated antibody at 1:15000 dilution for 1 h prior to color development. ABTS substrate was added for 15 min and subsequently quenched with 1% SDS prior to reading absorbance at 405 nm.

Results are shown in absorbance units (a.u.) as the mean value ± standard deviation (S.D.) on n = 3 technical replicates.

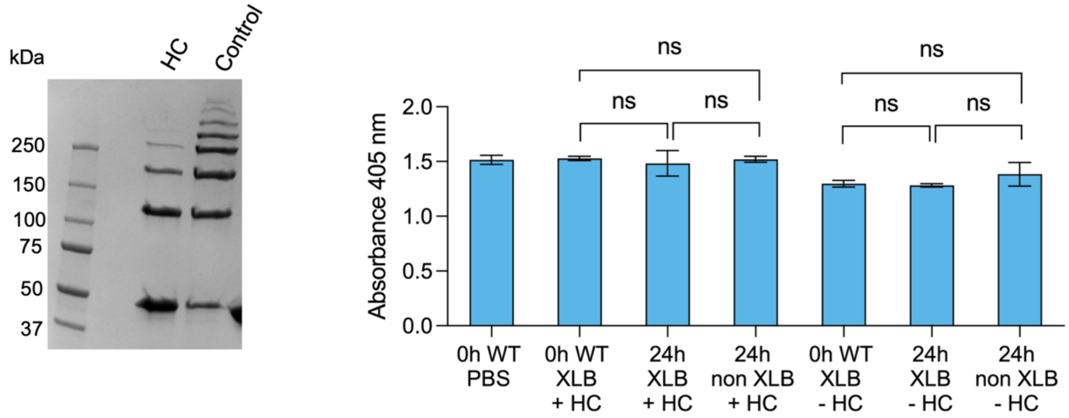

Fig. S8: Crosslinking with hexachlorophene

FlgE was treated in the presence or absence of 200 μM HC for 24 h prior to coating in (i) crosslinking buffer (XLB) (40 mM Tris pH 8.5, 160 mM NaCl, 1 M ammonium sulfate) with 1 mM NEM, or (ii) non-crosslinking buffer (non XLB) (40 mM Tris pH 8.5, 160 mM NaCl) with 1 mM NEM. The SDS-PAGE gel on the left shows the extent of inhibition by HC prior to coating. Buffer conditions were varied in the ELISA during coating to test sensitivity of the coating process to crosslinking buffer. HC was washed from the wells prior to adding the conjugate and performing the crosslinking reaction. All absorbance values obtained were comparable to that of control (0 h pre-incubation of coat protein, in PBS) indicating both that coating in crosslinking buffer did not affect the assay and that no inhibition occurred when HC was washed from the wells.

Results are shown in absorbance units (a.u.) as the mean value ± standard deviation (S.D.) on n = 3 technical replicates. Statistical significance was calculated using a 1-way ANOVA with post-hoc Tukey test ($p$ >0.05 ns, $p$ < 0.05*, 0.01** and 0.001***).

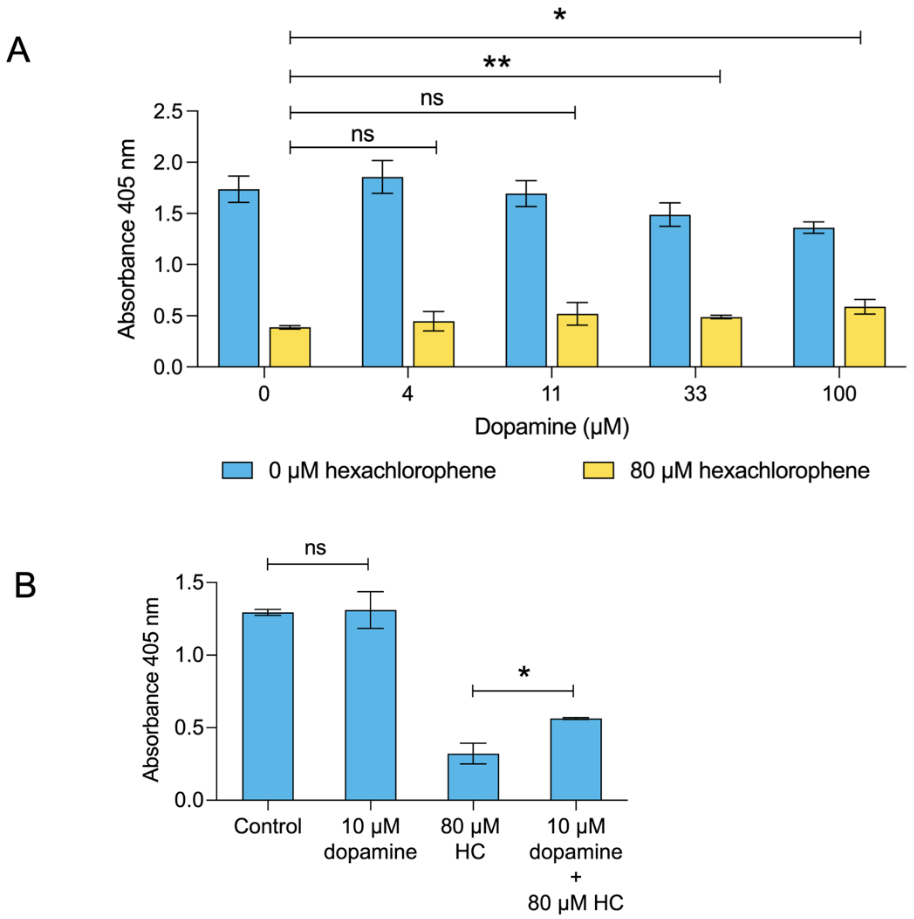

Fig. S9: Effects of dopamine

(A) Dopamine was added during the crosslinking reaction from 0-100 µM either alone or in the presence of 80 µM HC. Dopamine was identified as an activator using the NanoLuc assay. In the ELISA, dopamine alone at higher concentrations slightly lowered the signal. However, it slightly reverses the inhibition caused by HC.

(B) ELISA performed in the absence of NEM with dopamine and/or hexachlorophene. For the control, 1 mM NEM was used in the assay. Dopamine can generate signal comparable to control even in the absence of NEM. It also slightly reverses HC inhibition even at a low concentration.

Results are shown as the mean value ± standard deviation (S.D.) on n = 3 technical replicates. Statistical significance was calculated for pairwise comparisons using two-tailed t-tests ($p$ >0.05 ns, $p$ < 0.05*, 0.01** and 0.001***).

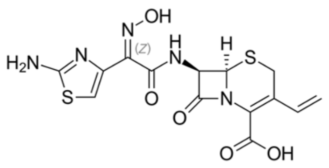

Cefdinir
214.4%

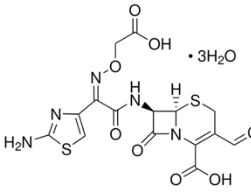

Cefixime trihydrate
195.3%

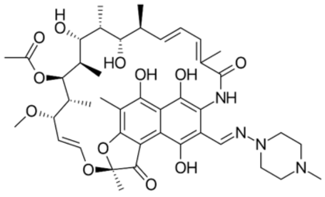

Rifampicin
186.5%

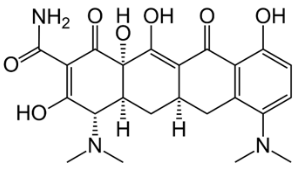

Minocycline hydrochloride
171.3%

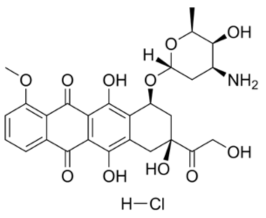

Doxorubicin hydrochloride
150.6%

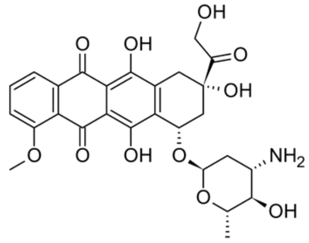

Epirubicin hydrochloride
146%

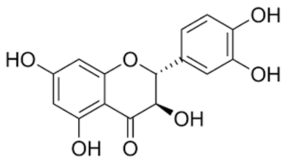

Taxifolin-(+)
144.6%

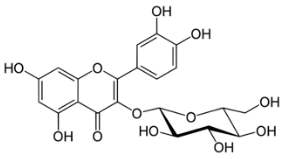

Isoquercitrin
138.3%

Fig. S10: Compounds identified during HTS that activated crosslinking by increasing the HRP-signal

Compounds are listed with the percentage of crosslinking over the no-compound control. These compounds group into specific classes - cephalosporins, tetracycline compounds and quercetins.

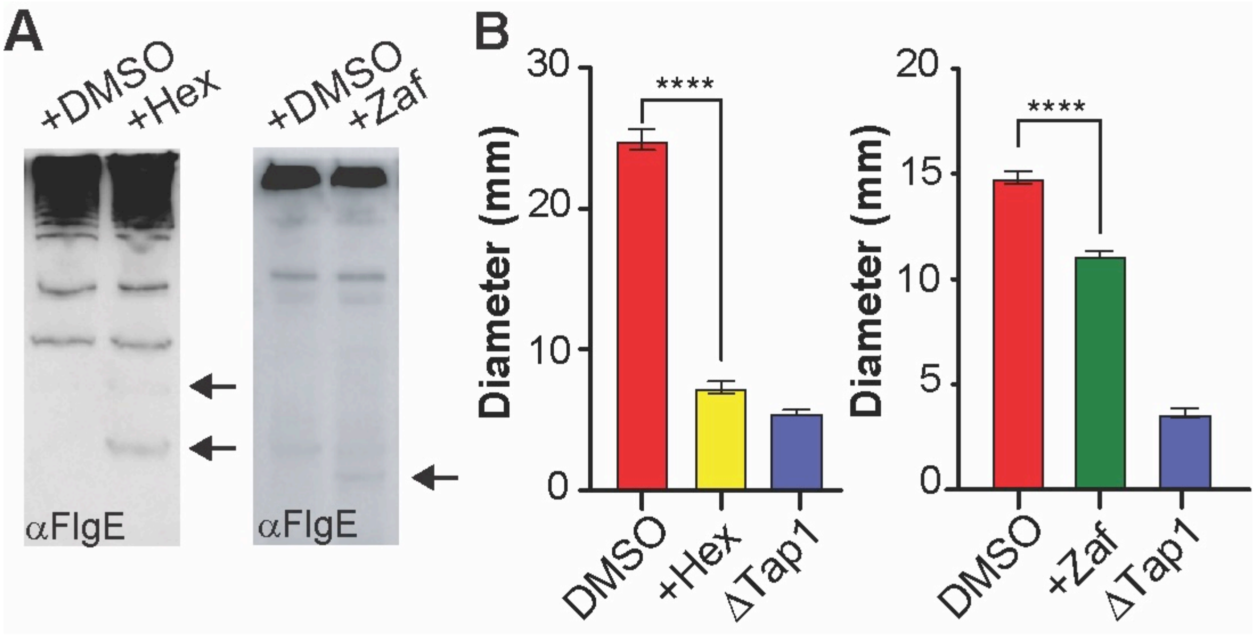

Fig. S11: Effects of zafirlukast on FlgE crosslinking and motility of *T. denticola* in comparison to the effects of hexachlorophene.

(A) αFlgE western blot (right) of *T. denticola* cultures treated with DMSO, hexachlorophene (83.3 µM, +Hex) or zafirlukast (50 µM, +Zaf). Black arrows denote the appearance of molecular weight bands that are lacking in DMSO-treated samples. B) Swimming plate assays of WT, ΔTap1, Hex-treated and Zaf-treated *T. denticola* cells at the same concentrations as those used in (A). Swimming plate assays were performed in triplicate and the diameter of each strain reported as the average diameter ± the standard deviation of three biological replicates. Statistical significance was calculated using a two-tailed Student's t-test ($p<0.05$*, 0.01** and 0.001****). Hexachlorophene data is taken from[1] to show in comparison.
